# Novel fat-taste enhancers modulate functional connectivity of the reward system following sustained activation of gustatory pathways in mice

**DOI:** 10.1101/2025.09.11.675492

**Authors:** Lidia Cabeza, Lucas Jantzen, Christophe Houdayer, Emmanuel Haffen, Aziz Hichami, Naim Akhtar Khan, Vincent Van Waes

**Author notes:** **Corresponding author** Lidia CABEZA - Mail address: 19, Rue Ambroise Paré, 25030, Besançon Cedex, France.

## Abstract

Humans and rodents exhibit an innate preference for dietary fat, and dysfunction of lingual fat-taste receptors CD36 and GPR120 in obesity suggests that impaired oro-sensory lipid detection may contribute to its development. Rodent models with reduced fat-taste sensitivity display increased fat intake, highlighting the role of taste perception in dietary regulation. The newly developed NKS-3 and NKS-5, high-affinity agonists for CD36 and CD36/GPR120 receptors, respectively, induce early fat satiation and reduce both fat-rich food intake and body weight gain in diet-induced obese mice. However, the neural mechanisms underlying these effects remain unclear. Here, we investigated the response of the reward system to a sustained activation of CD36 and GPR120 via fat-taste enhancers NKS-3 and NKS-5 compared to linoleic acid (LA). Male C57BL/6JRj mice (N=88) received oral administration of vehicle, 0.2% LA, 50 µM NKS-3 or 75 µM NKS-5 for 10 days. The immunohistochemical analysis of cerebral FosB neuronal expression revealed a reorganization of the functional network connectivity. Behavioral assessments over an additional 10-day period in a second cohort of animals detected no adverse motivational, compulsive-like, depressive or anxiogenic effect of the treatments. Our findings suggest that our fat-taste enhancers may offer a promising therapeutic strategy against obesity, leveraging the oral-gut-brain axis to regulate fat-rich food intake.

## Introduction

The innate preference for dietary fat in human beings and rodents is primarily regulated by fat-taste receptors like CD36 and G protein coupled receptor 120 (GPR120), located in taste bud cells and, respectively, involved in early detection and post-ingestive regulation of dietary fat intake [1–4]. Former studies have shown that dietary fat is able to activate several brain regions [5,6]. In a preclinical study, we have previously demonstrated that oral lipid taste perception triggers the activation of canonical and reward pathways [7]. Indeed, lingual fat-taste receptors trigger signaling events that convey to the first relay of peripheral information in the brain stem, the nucleus of the solitary tract (NST), via the facial chorda tympani (VII) and glossopharyngeal (IX) cranial nerves. The gustatory information is sent to the parabrachial nucleus (PBN) and then reaches the gustatory ventral posteromedial thalamic nucleus (VPM), to continue up to the gustatory insular cortex. This gustatory cortex integrates information on taste qualities and receives modulatory input from the central amygdala (CeA) and posterior areas of the lateral hypothalamus, including the parasubthalamic nucleus (PSTN). The reward system, including the ventral tegmental area (VTA), responds to the flow of this information and contributes to reshape feeding behavior [8].

The expression of CD36 and GPR120 receptors at peripheral, e.g. small intestine, and central level, e.g. hypothalamus [9,10], endorses the construct of “functional continuum” along the oro-intestinal-brain axis in lipid sensing. Accordingly, defective fat perception along this continuum may affect overall feeding behavior, including food preference and satiety. Supporting this idea, the expression of lingual fat-taste receptors has been found downregulated in most patients suffering from obesity [11–13]. Moreover, increased fat intake has been reported in murine models due to reduced fat-taste sensitivity [3], which also points to the role of taste perception in eating behavior.

In this context, we have developed high-affinity agonists for CD36 and GPR120 receptors, named NKS-3 and NKS-5 [14], in order to study their potential as an alternative therapeutic strategy in obesity. Through *in vivo* modulating fat perception, these synthetic molecules derived from linoleic acid (LA), were found to induce early satiation and reduce both fat-rich food intake and body weight gain in diet-induced obese mice. In addition to other beneficial effects, e.g. induced insulin sensitivity, normalized dyslipidemia [14], we have recently demonstrated that NKS-3 is able to significantly reduce systemic and central inflammation [15]. However, the neural mechanisms underlying these effects, and in particular the effects on feeding behavior, remain to be elucidated. Hence, it was thought worthwhile to study the behavioral and neural response to sustained stimulation of CD36 and GPR120 via fat-taste enhancers NKS-3 and NKS-5 in naïve adult mice. Understanding the mechanisms of action of these new pharmacological tools is essential to foresee an interventional approach in pathological conditions as obesity.

## Materials and methods

### Animals

Eighty-eight 6-to-8 week-old male C57BL/6JRj mice (*Ets Janvier Labs*, Saint-Berthevin, France) were used in this study. Animals were group-housed and maintained under a controlled environment (12-h light/dark cycle; temperature: 22±2 °C; humidity: 55±10%). They had *ad libitum* access to standard food (KlibaNafag3430PMS10, *Serlab*, CH-4303 Kaiserau, Germany) and vehicle or treatment during the entire duration of the experiment, and their weight was regularly monitored.

All experimental procedures were conducted in accordance with the Guide for the Care and Use of Laboratory Animals (NIH), the Animal Research: Reporting of In Vivo Experiments (ARRIVE) guidelines, and the European Union regulations (Directive 2010/63/EU). The procedures were approved by the Université Marie et Louis Pasteur Animal Care and Use Committee (CEBEA-58).

### Pharmacological treatments

After a 6-day acclimation period to the animal facilities, mice were divided into four experimental groups comprising 22 individuals each: a control group treated with vehicle (VEH; 0.3% w/v Arabic gum solution, in water; G9752, *Sigma-Aldrich*, France), a LA-treated group (0.2% LA w/v, in VEH; L1376, *Sigma-Aldrich*, France), a NKS-3-treated group (diethyl (9Z,12Z)-octadeca-9,12-dien-1-ylphosphate (C22H43PO3), 50 µM, in VEH) and a NKS-5-treated group (9Z-12Z-methyloxetan-3-yl)methyl octadeca-9,12-dienoate (C23H40O3), 75 µM, in VEH). The NKS-3 and NKS-5 agonists were produced and conserved as previously described [14] and treatments were freshly prepared twice a week in order to minimize lipid oxidation. Beverage and food intake were carefully monitored.

A subset of individuals representative of all experimental conditions (Group A, n=48) were treated for 12 days before they performed a battery of behavioral tests aiming at evaluating the effect of the molecules on their emotionality. The remaining animals (Group B, n=40) were euthanized after 10 days of pharmacological treatment: a subset of 28 brains randomly selected were collected and processed for immunostaining analysis (see **Figure 1A** for the timeline of the experimental design).

**Figure 1.**
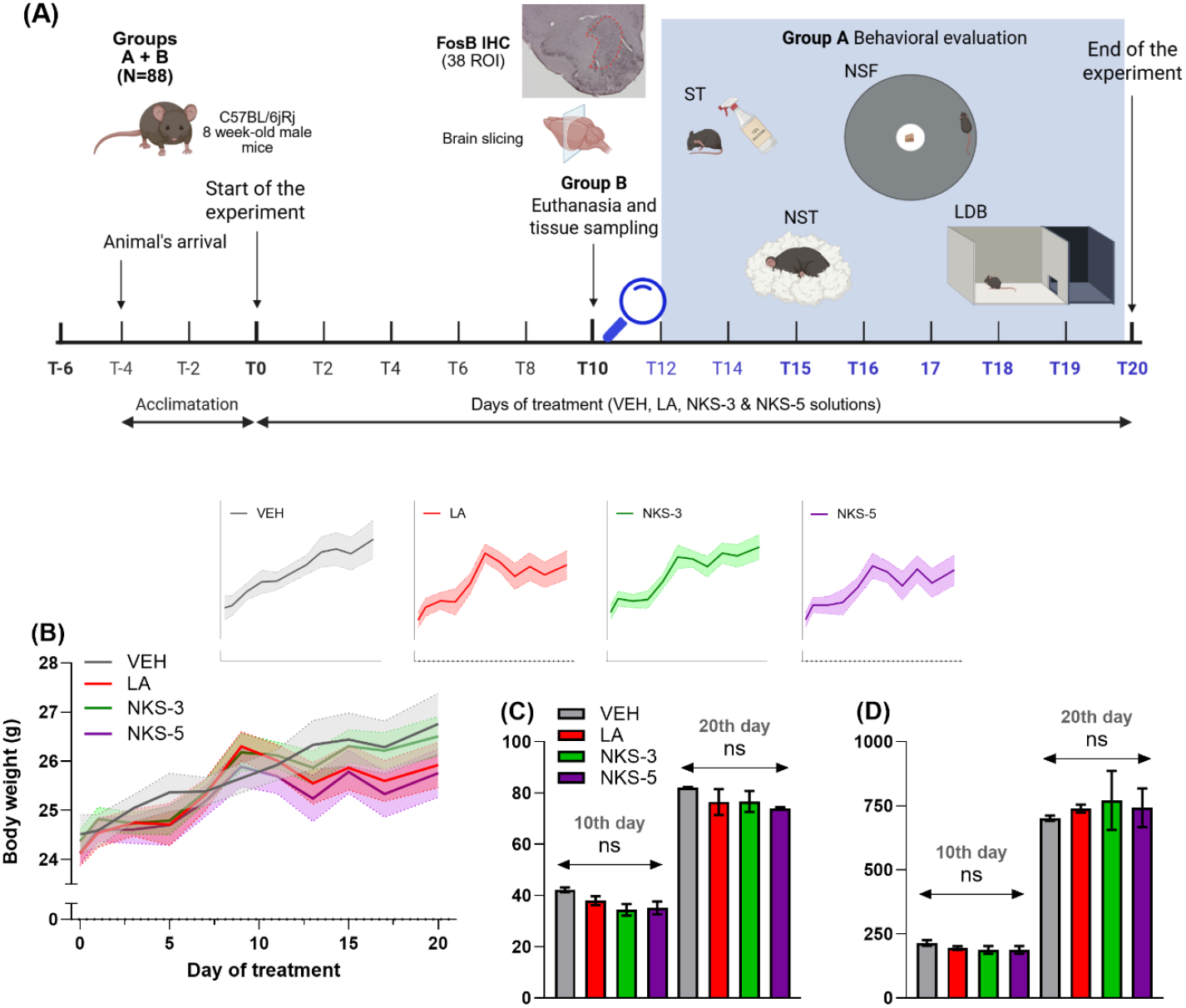
Experimental design and mice monitoring. **(A)** After 6 days of acclimatation to the animal facilities, animals (Groups A+B) were treated with either vehicle (VEH), 0.2% linoleic acid (LA), 50µM NKS-3, or 75µM NKS-5 solutions. Mice from the Group B (n=40) were euthanized after 10 days of treatment, and a subset of brains (n=28) were collected and processed for immunostaining targeting FosB expression. Animals from the Group A continued being treated and were emotionally evaluated via a battery of behavioral tests from the 12^th^ day (created with *BioRender*.*com*). **(B)** All animals gained weight during the experiment and no effect of the treatments was observed in their progression (Group A+B, mean weight at 10^th^ day (g) ± SEM, VEH: 25.86±0.30; LA: 25.86±0.30, NKS-3: 26.06±0.25, NKS-5: 25.62±0.35; Group A at 20^th^ day, VEH: 26.76±0.62; LA: 25.93±0.46, NKS-3: 26.51±0.41, NKS-5: 25.76±0.49). **(C)** Mice from the four experimental conditions consumed similar amounts of food (cumulated food intake at 10^th^ day (g) ± SEM, VEH: 42.25±0.88; LA: 38.04±1.77; NKS-3: 34.49±2.25; NKS-5: 35.22±2.48; at 20^th^ day, VEH: 82.05±0.33; LA: 76.51±5.06; NKS-3: 76.70±4.06; NKS-5: 73.95±0.48), and **(D)** took similar volumes of VEH or treatments, regardless the time point evaluated (cumulated beverage intake at 10^th^ day (g) ± SEM, VEH: 215.05±11.18; LA: 195.10±7.40; NKS-3: 187.53±15.42; NKS-5: 187.57±15.42; at 20^th^ day, VEH: 701.87±10.55; LA: 739.44±15.35; NKS-3: 771.29±115.13; NKS-5: 742.68±75.44). ns, not significant.

### Behavioral characterization

The behavioral evaluation of the animals from the Group A was conducted during the light/inactive phase of the circadian cycle (starting at 8 a.m.) following operative protocols from our laboratory [16,17]. Depressive-like behavior was assessed using the splash test (ST), through leveraging mice’s inherent drive to groom themselves [18]. Scores from the nestlet shredding test (NST) informed about the emergence of repetitive, compulsive-like behavior [19,20]. Latencies to consume food in a novel environment (i.e. hyponeophagia), after food deprivation, informed of appetitive incentive motivation in the novelty suppressed feeding task (NSF) [21,22]. Finally, anxious-like behavior was evaluated in the light-dark box test (LDB) by presenting animals with a choice paradigm, where they faced an internal conflict between staying in a dimly lit ‘safe’ compartment and moving into a brightly lit ‘novel’ compartment [23]. A 24 h washout period was included between consecutive behavioral tests in order to limit stress effects, with the exception of the NSF task, which required a 48 h washout period due to food deprivation.

### Animals sacrifice and tissue sampling

After 10 days of pharmacological treatment, animals from the Group B were euthanized for evaluation of sustained neural activation by FosB immunostaining as we previously described [17,24]. In brief, after an intraperitoneal injection of pentobarbital (55 mg/kg, Exagon®, *Med’Vet*, France), mice were transcardially perfused with ice-cold 0.9% NaCl, followed by 4% paraformaldehyde (PFA, *Roth®*, Karlsruhe, Germany) dissolved in 0.1 M phosphate buffer (PB, pH 7.4). Collected brains were post-fixed overnight at 4 °C in PFA and then cryoprotected for 24 h by immersion in an ice-cold 15% sucrose solution (D(+)-Saccharose, *Roth®*, Karlsruhe, Germany) in a 0.1 M PB. Brains were finally frozen by immersion in isopentane (2-methylbutane, *Roth®*, Karlsruhe, Germany) and sliced into 30-µm-thick coronal serial sections.

### FosB immunohistochemistry

FosB immunostaining was conducted across 38-targeted regions either intrinsic to or involved in the modulation of the brain’s reward system, encompassing ten distinct coronal sections. See detailed information of regions and Bregma coordinates in **Table 1**.

**Table 1.**
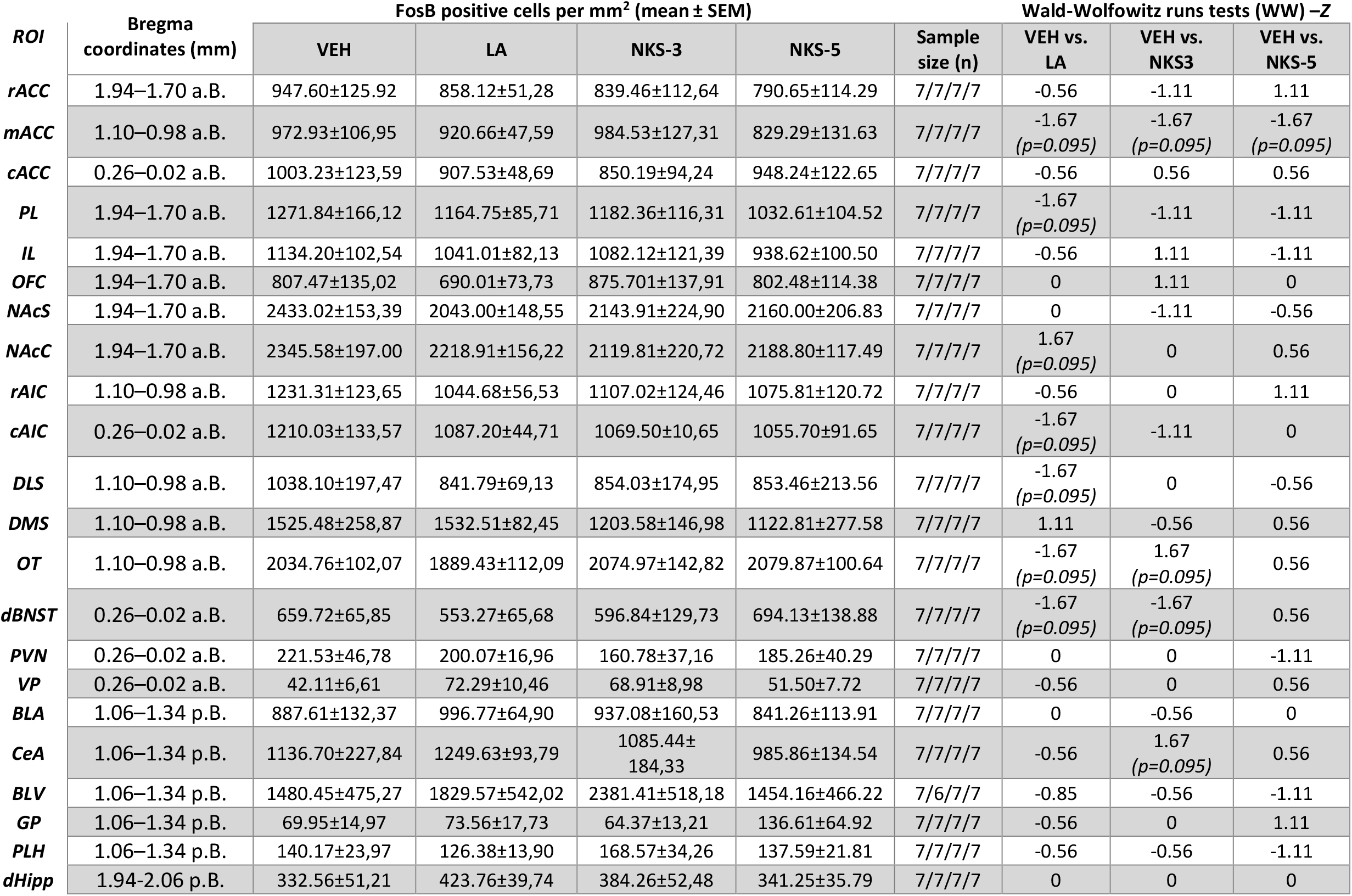
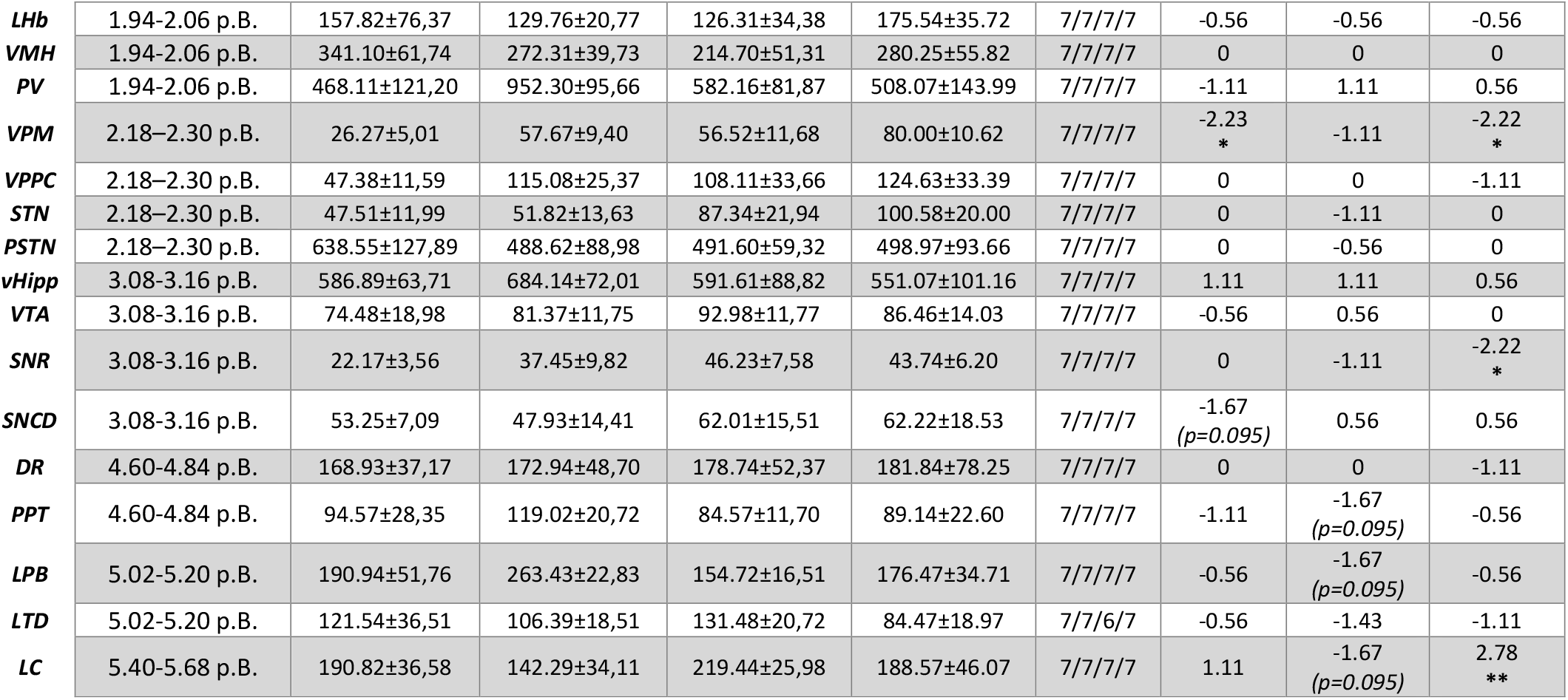
FosB positive cells density (sustained neuronal activity) and statistical comparisons between experimental conditions and brain regions of interest (ROI). Bregma coordinates (mm) for sections selection: anterior to Bregma – aB; posterior to Bregma – pB. Treatment: vehicle – VEH; linoleic acid –LA; fat-taste enhancers NKS-3 and NKS-5. Data are means ± SEM. Rostral anterior cingulate cortex – rACC, medial anterior cingulate cortex – mACC, caudal anterior cingulate cortex – cACC, prelimbic cortex – PL, infralimbic cortex – IL, orbitofrontal cortex – OFC, shell of the nucleus accumbens – NAcS, core of the nucleus accumbens – NAcC, rostral anterior insular cortex – rAIC, caudal anterior insular cortex – cAIC, dorsolateral striatum – DLS, dorsomedial striatum – DMS, olfactory tubercle – OT, dorsal region of the bed nucleus of the stria terminalis – dBNST, paraventricular nucleus of the hypothalamus – PVN, ventral pallidum – PV, basolateral amygdala – BLA, central amygdala – CeA, basolateral ventral nucleous of the amygdala – BLV, globus pallidus – GP, peduncular part of the lateral hypothalamus –PLH, dorsal hippocampus – dHipp, lateral habenula – LHb, ventromedial hypothalamus – VMH, paraventricular thalamic nucleus – PV, ventral posteromedial thalamic nucleus – VPM, parvicellular part of the ventral posterior thalamic nucleus – VPPC, subthalamic nucleus of the hypothalamus – STN, parasubthalamic nucleus of the hypothalamus – PSTN, ventral hippocampus – vHipp, ventral tegmental area – VTA, subtantia nigra pars reticulate – SNR, dorsal substantia nigra compacta – SNCD, dorsal raphe – DR, pedunculopontine nucleus – PPT, lateral parabrachial nucleus – LPB, laterodorsal tegmental nucleus – LDT, and nucleus coeruleus – LC, according to Franklin and Praxinos [32]. n: sample size. *, p<0.05; **, p<0.01.

After blocking endogenous peroxidase activity (0.3% hydrogen peroxide solution, 20 min), floating sections were incubated with the primary antibody (1:1000; ab184938, rabbit anti-FosB, *Abcam*) for 48 h, and then with the secondary HRP antibody (1:1000; BA-1000, goat anti-rabbit Ig, *Vector Laboratories*) for 24 h, both at 4 °C. An avidin horseradish peroxidase complex (ABC Elite kit, *Vector Laboratories*) served to amplify the signal (40 min at room temperature), and a 3,3′-Diaminobenzidine (DAB) chromogen solution to visualize the peroxidase complex. Sections were then mounted on gelatin-coated slides, dehydrated with successive alcohol baths (70°, 95°, 100°, and 100°, 3 min), cleared with xylene (*Avantor*^®^, 3×5 min) and cover-slipped with Canada Balsam (*Roth*^®^, Karlsruhe, Germany). Photomicrographs for analysis were acquired using a 4x objective of an Olympus microscope Bx51 equipped with an Olympus DP50 camera. *ImageJ* software (National Institute of Health, Bethesda, Maryland, USA, http://imagej.net/ij/ [25] was used to quantify FosB-labelled cell nuclei within frames manually traced and delimiting the brain region of interest (ROI). See **Table 1** for detailed final samples sizes.

### Data and statistical analyses

The results are presented as means ± SEM. The statistical analyses were conducted using STATISTICA 10 (*Statsoft*, Palo Alto, United States) and *R* software (version 4.4.1; R Core Team, 2024; https://www.R-project.org/). Figures were designed using *GraphPad Prism* version 10.5.0.774 for Windows (*GraphPad* software, Boston, Massachusetts USA, "https://www.graphpad.com), and networks diagrams were constructed using *R* analysis (*igraph* package) and illustration (*Cytoscape* software, version 3.10.3) tools.

Assumptions for parametric analysis were verified using Shapiro-Wilk and Levene’s tests. The progression of the body weight was evaluated with a repeated-measures ANOVA (RMA) design, including four groups (VEH, LA, NKS-3 and NKS-5) as between-subject factor (treatment), and 10 or 20 measurements as within-subject factor (time in days). Cumulated food and beverage intake were compared at 10 and 20 days of treatment using a One-way ANOVA. When the data sets did not meet assumptions for parametrical analysis, Kruskal-Wallis (KW) with *post hoc* Dunn’s correction, and Wald-Wolfowitz runs tests (WW) were used. The NSF score curves were compared using the Gehan-Breslow Wilcoxon (GBW) test. Finally, Pearson *r* values from interregional FosB expression data were correlated using cross-correlation matrices within each experimental condition. After applying a threshold of 0.7 (allowing to eliminate weak associations, adapted for the study of the network’s core), the similarity of the cross-correlation matrices was studied using the RV (*rhô-vectoriel*) coefficient method followed by a permutation test. Network diagrams were topologically compared using different statistical analysis: (i) the interconnection between the identified nodes was studied through a clustering coefficient; (ii) the connectivity extent was studied through the measure of the degree of the nodes’ centrality; (iii) the efficiency of the network’s connections was approximated through the method of the shortest path length; (iv) and the organization of the networks was compared using modularity analyses. The threshold for statistical significance was set at p<0.05 and dependencies with p≤0.1 were considered as trends.

## Results

### Body weight and consummatory behavior

Continuous monitoring of the animals showed an expected increase in body weight over time, without any significant effect of the treatments [RMA, Group A+B: main effect of treatment, F_3,84_=0.3, p=0.84; mean effect of time, F_3,252_=278.4, p<0.0001; treatment × time interaction, F_9,252_=1.8, p=0.06; Group A: treatment, F_3,44_=0.9, p=0.46; time, F_4,176_=30.5, p<0.0001; treatment × time interaction, F_1,176_=0.9, p=0.55] (**Figure 1B**). The consummatory behavior of mice remained unaltered, i.e., no effect of the treatments was observed either on food or beverage intake [cumulated food intake at 10^th^ day, Group A+B: H_3,16_=6.3, p=0.096; at 20^th^ day, Group A: H_3,8_=4.0, p=0.26; cumulated beverage intake at 10^th^ day, Group A+B: H_3,16_=3.4, p=0.33; at 20^th^ day, Group A: H_3,8_=0.7, p=0.88] (**Figure 1C,D**).

### Behavioral evaluation

Our results indicate no depressive, anxiogenic, or compulsive-like effects of the treatments on the behavior of mice from the group A after 12 days of treatment. Animals from the four experimental conditions spent similar amounts of time grooming themselves in the ST [H_3,24_=2.2, p=0.53] (**Figure 2A**). No effect of the treatments was observed on mice’s shredding activity in the NST [H_3,48_=4.9, p=0.18] (**Figure 2B**). Similarly, the treatments did not impact the behavior of mice during the NSF task, as shown by the similar latencies to start consuming the presented food [GBW: X^2^=4.3, p=0.23] (**Figure 2C**). Finally, the behavior of mice treated either with LA, NKS-3 or NKS-5 solutions did not differ from the control group in the LDB [entries into lit compartment, H_3,48_=2.3, p=0.51; time in lit compartment, H_3,48_=1.2, p=0.74; time in transition zone, H_3,48_=0.5, p=0.93] (**Figure 2D-F**), demonstrating the absence of anxiogenic effects. However, a significant difference in the motor activity between LA- and NKS-5-treated animals was observed [H_3,48_=8.7, p=0.03; *post hoc* Dunn’s test, LA *vs*. NKS-5: Z=2.8, p=0.03] (**Figure 2G**).

**Figure 2.**
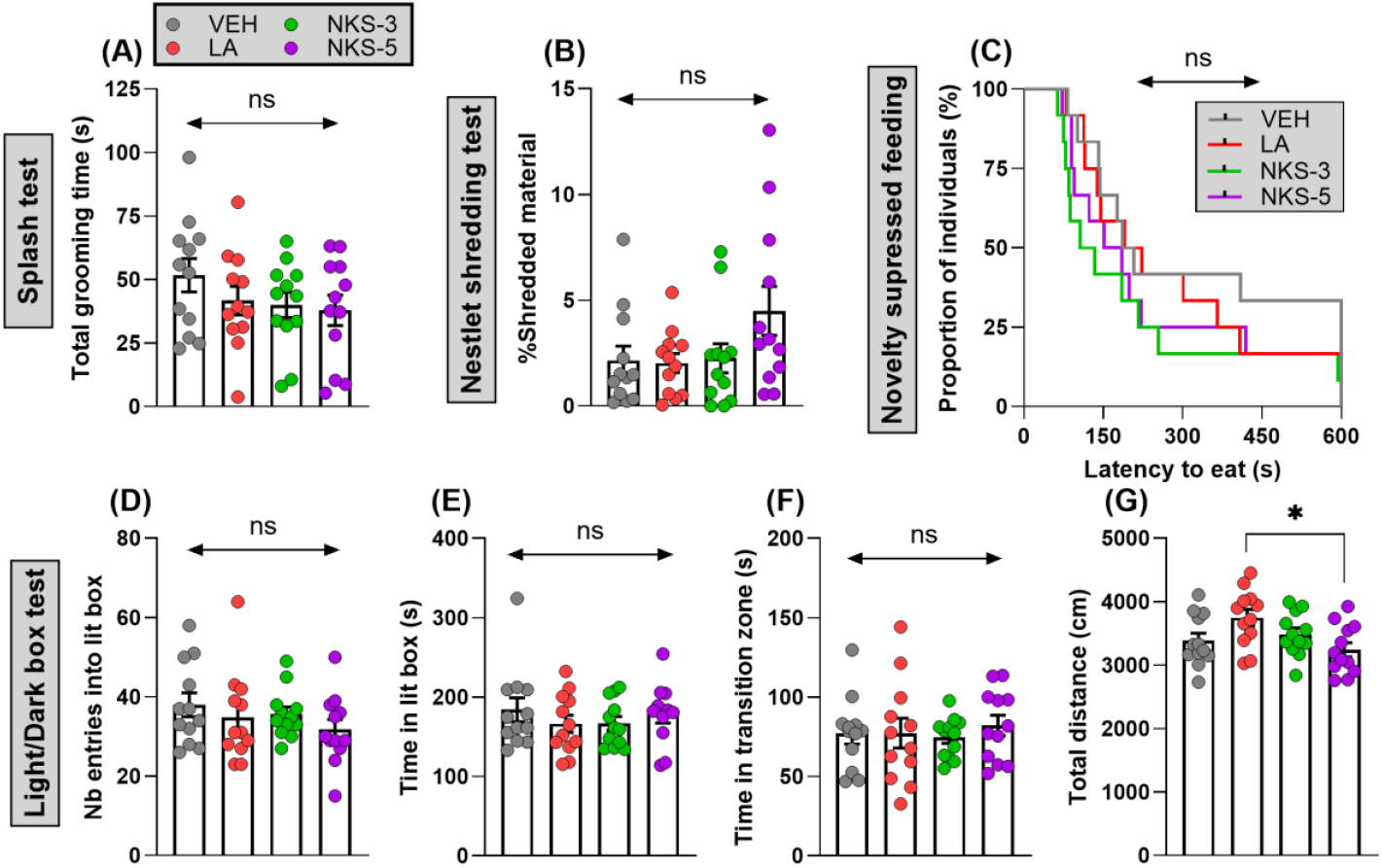
Effect of linoleic acid (LA), NKS-3 and NKS-5 exposure on emotional behavior. **(A)** The total time of grooming (mean grooming time (s) ± SEM; VEH: 51.65±6.58; LA: 41.68±5.64; NKS-3: 40.01±5.04; NKS-5: 37.87±5.99) in the splash test (ST) was similar between the animals of the four experimental groups. **(B)** The treatments did not induce the emergence of repetitive, compulsive-like behavior in mice, as evidenced in the nestlet shredding test (NST) (shredded material (%) ± SEM; VEH: 2.15±0.67; LA: 2.03±0.45; NKS-3: 2.25±0.69; NKS-5: 4.49±1.16). **(C)** All animals exhibited similar latencies to eat in the novelty-suppressed feeding (NSF) task, attesting the absence of motivational deficits among the experimental groups (latency to eat (s) ± SEM; VEH: 320.60±63.98; LA: 273.30±53.15; NKS-3: 206.30±55.45; NKS-5: 236.90±55.77). **(D-F)** The light-dark box (LDB) test did not evidence an anxiogenic effect of the treatments, as shown by the similar values obtained for the different scores evaluated (entries into lit compartment ± SEM; VEH: 38.00±2.99; LA: 34.83±3.31; NKS-3: 35.67±1.78; NKS-5: 31.75±2.53; time in lit compartment ± SEM; VEH: 184.00±14.94; LA: 166.30±10.89; NKS-3: 166.70±8.53; NKS-5: 178.40±10.92; time in transition zone (s) ± SEM; VEH: 77.06±6.67; LA: 77.23±9.38; NKS-3: 74.48±3.69; NKS-5: 82.13±6.42). **(G)** However, an isolated significant difference was observed in the total distance travelled during the LDB test, with LA-treated mice exploring more than NKS-5-treated mice (mean distance (cm) ± SEM: VEH: 3387±116.30; LA: 3750±128.70; NKS-3: 3487±97.33; NKS-5: 3243±112.00). *, p<0.05; ns, not significant.

### Sustained neural activation through FosB expression

The levels of sustained neuronal activation addressed through FosB expression in the brain ROIs, as well as the statistical comparisons between experimental conditions, are presented in **Table 1**.

Among the targeted brain areas, the density of FosB-labelled cells was significantly increased in the VPM of LA-treated mice [WW, VEH *vs*. LA: Z=-2.2, p=0.026] and of NKS-5-treated mice [VEH *vs*. NKS-5: Z=-2.2, p=0.026]. Compared to control individuals, NKS-5-treated mice also presented a significant increase of FosB positive density in the *substantia nigra pars reticulata* (SNR) [VEH *vs*. NKS-5: Z=-2.2, p=0.026], and a significant decrease in the *locus coeruleus* (LC) [VEH *vs*. NKS-5: Z=2.8, p=0.0054]. The density of FosB-labelled neurons in LA-treated mice tended to be significantly reduced in the medial anterior cingulate cortex (mACC), the prelimbic cortex (PL), the core of the nucleus accumbens (NAcC), the caudal part of the anterior insular cortex (cAIC), the dorsolateral striatum (DLS), the paraventricular hypothalamic nucleus (PVN) and the dorsal *substantia nigra pars compacta*; and tended to be significantly increased in the dorsomedial striatum (DMS) [all ps<0.1]. Compared to the control condition, the FosB density of NKS-3-treated individuals tended to be significantly reduced at the level of the mACC, the dorsal part of the bed nucleus of the *stria terminalis* (dBNST), the CeA, the pedunculopontine nucleus (PPT) and the LPB; and tended to be significantly increased in the olfactory tubercle (OT) and the LC [all ps<0.1]. Finally, in NKS-5-treated mice, the density of FosB-labelled cells was found to tend to a significant reduction only at the level of the mACC [p<0.1].

The previous findings led us to study functional connectivity by computing covariance across the experimental groups on FosB expression aiming at assessing the effect of the treatments on our reward network’s interactions (**Figure 3A-D**). The analysis by the RV coefficient method indicates that the cross-correlation matrix for the LA condition differs from those of control (VEH *vs*. LA, RV: 0.32), NKS-3 (NKS-3 *vs*. LA, RV: 0.34) and NKS-5 (NKS-5 *vs*. LA, RV: 0.35) conditions. While NKS-3 and NKS-5 cross-correlation matrices share a moderate similarity (NKS-3 *vs*. NKS-5, RV: 0.65), they have a close correlation structure to the VEH matrix (VEH *vs*. NKS-3, RV: 0.72; VEH *vs*. NKS-5, RV: 0.77). The permutation analysis confirmed a shared common structure between the experimental conditions.

**Figure 3.**
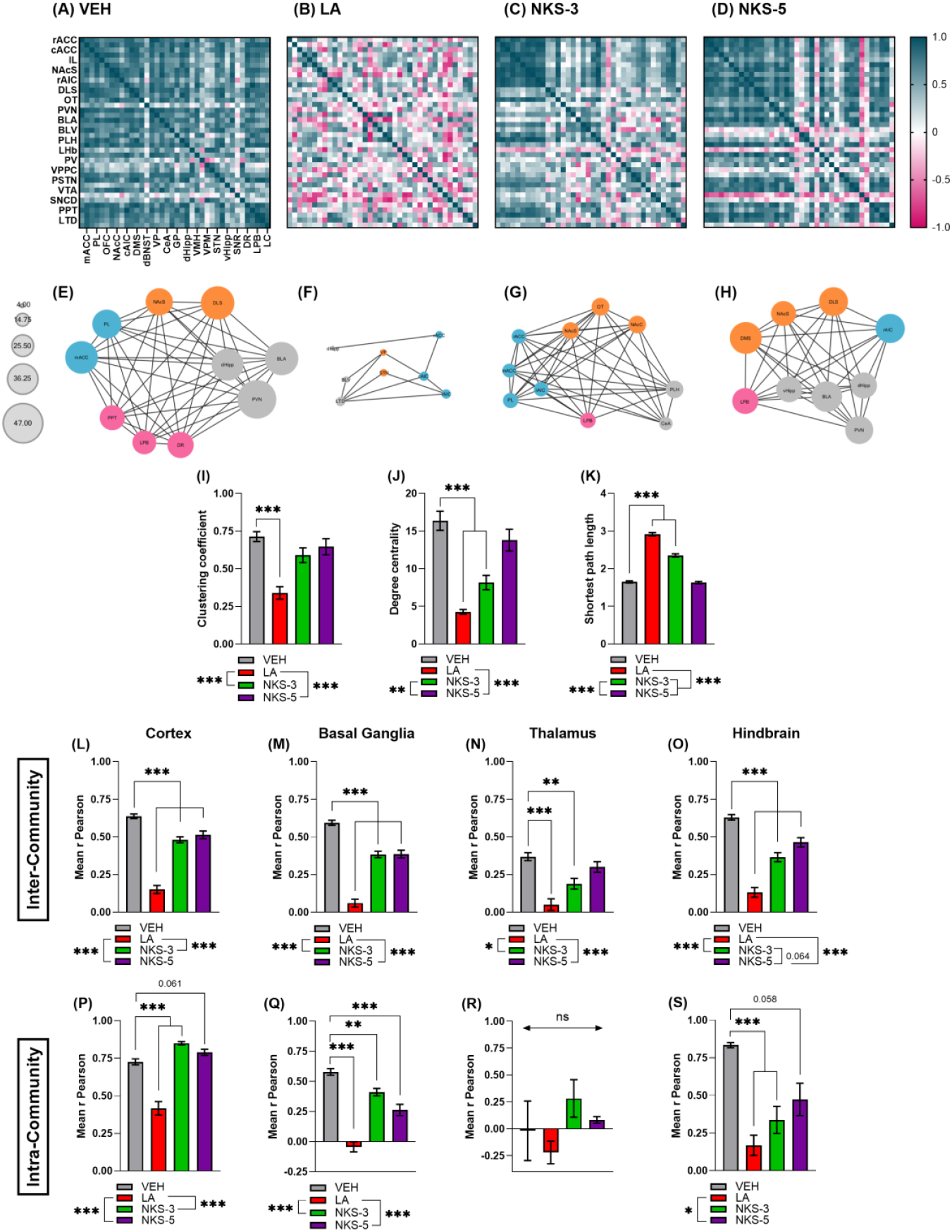
Generation of reward system’s neuronal activity networks. **(A-D)** Cross-correlations matrices showing interregional correlations for FosB expression after 10 days of vehicle (VEH), linoleic acid (LA), NKS-3 or NKS-5 treatment. Brain structures are organized rostro-caudally. Colors illustrate correlation strength based on Pearson’s r (blue, positive correlation; pink, negative correlation; white, no correlation). **(E-H)** Networks were generated by significant correlations and illustrate the strongest 25% of them. The diameter of the nodes reflects their importance, i.e. the degree of their connectivity (cortex, blue nodes; basal ganglia, orange nodes; hindbrain, pink nodes; other regions, grey nodes). **(I)** The mean clustering coefficient of the system was significantly reduced by LA, but not affected by NKS-3 and NKS-5 administration (VEH: 0.71±0.03; LA: 0.34±0.04; NKS-3: 0.59±0.05; NKS-5: 0.65±0.05). **(J)** The connectivity of the network’s nodes was reduced after LA and NKS-3 administration, and no effect of NKS-5 was observed (mean degree centrality ± SEM, VEH: 16.37±1.28; LA: 4.26±0.31; NKS-3: 8.16±0.95; NKS-5: 13.79±1.45). **(K)** The increased shortest path length in LA and NKS-3 conditions indicate weaker and less direct connections within the network compared to the control condition (mean distance between nodes ± SEM, VEH: 1.65±0.03; LA: 2.92±0.04; NKS-3: 2.35±0.04; NKS-5: 1.63±0.03). **(L-S)** Mean *r* scores were calculated from interregional correlation coefficients of four different communities of brain regions (cortex, basal ganglia, thalamus and hindbrain), between them and the rest of the network –inter-community scores, and between themselves –within-community scores (cortex, VEH-inter: 0.64±0.02; VEH-within: 0.73±0.02; LA-inter: 0.15±0.03; LA-within: 0.42±0.04; NKS-3-inter: 0.48±0.02; NKS-3-within: 0.85±0.01; NKS-5-inter: 0.51±0.03; (NKS-5-within: 0.79±0.02; basal ganglia: VEH-inter: 0.59±0.02; VEH-within: 0.58±0.03; LA-inter: 0.06±0.03; LA-within: −0.04±0.04; NKS-3-inter: 0.38±0.02; NKS-3-within: 0.41±0.03; NKS-5-inter: 0.39±0.03; NKS-5-within: 0.26±0.05; thalamus: VEH-inter: 0.37±0.03; VEH-within: −0.02±0.28; LA-inter: 0.05±0.04; LA-within: - 0.22±0.11; NKS-3-inter: 0.19±0.04; NKS-3-within: 0.28±0.17; NKS-5-inter: 0.30±0.23; NKS-5-within: 0.08±0.03; hindbrain, VEH-inter: 0.63±0.02; VEH-within: 0.83±0.02; LA-inter: 0.13±0.03; LA-within: 0.17±0.07; NKS-3-inter: 0.36±0.03; NKS-3-within: 0.34±0.09; NKS-5-inter: 0.46±0.03; NKS-5-within: 0.47±0.11). *, p<0.05; **, p<0.01; ***, p<0.001; ns, not significant.

Based on these results, we generated network diagrams where nodes represent individual brain regions, and links between pairs of nodes represent the strongest 25% of the calculated correlations (**Figure 3E-H**). Different network densities were observed at a global level (**Figure 3I**). Control animals show a well-preserved connectivity within the network, with dense and organized interactions. Mice treated with LA presented a more fragmented network, with a loss of connections compared to the control group, indicating a functional disorganization. The NKS-3 treatment induced a moderate modification of the connections that implies an average stability. The global network density of NKS-5-treated animals is similar to the control condition [F_3,148_=13.2, p<0.0001; *post hoc*, VEH *vs*. LA, p<0.0001; VEH *vs*. NKS-3, p=0.22; VEH *vs*. NKS-5, p=0.71]. These findings were confirmed by modularity analyses (modularity score, VEH: 0.16; LA: 0.50; NKS-3: 0.28; NKS-5: 0.12). However, different nodes, i.e. different brain regions, contribute to the functional networks (**Figure 3E-H**). The LA condition shares only one node with the VEH condition (dorsal hippocampus –dHipp), while the NKS-3 and NKS-5 conditions share four (PL, mACC, shell of the nucleus accumbens –NAcS, and LPB) and six (NAcS, DLS, PVN, dHipp, basolateral amygdala –BLA, and LPB) nodes respectively. By applying graph theoretical analysis to our networks, we addressed the relevance of these treatment-induced brain regions changes. First, we examined how the relative ranking of the degree centrality (i.e. number of links associated with a given node) differed in our networks. Compared to the control condition, the LA condition shows a strong loss of connections within a fragmented network [F_3,148_=25.2, p<0.0001; *post hoc*, VEH *vs*. LA, p<0.0001], while the reduction of connections in the NKS-3 condition suggests a partial, less severe fragmentation [p<0.0001]. The nodes’ connectivity observed in the NKS-5 condition was similar to the VEH condition [p=0.34] (**Figure 3J**). Secondly, we calculated the shortest path lengths of our networks, and evidenced the existence of weaker or more indirect connections in the LA condition compared to the control condition. The connections’ distance was found moderately increased in the NKS-3 condition, while that of the NKS-5 condition was similar to the control condition (**Figure 3J**).

Aiming at identifying further patterns, we selected four different sets of structures (cortex, basal ganglia, thalamus and hindbrain) and used them to (i) compute participation coefficients measuring how connected a community is to the rest of the network; and to (ii) compute within-community comparisons of mean correlation scores in order to approximate how well connected brain regions are to its own community. Cortical regions display both high inter- and within-community scores in the control condition, which were significantly decreased with the LA treatment [VEH *vs*. LA-inter, F_3,956_=86.2, p<0.0001; *post hoc*, p<0.0001; VEH *vs*. LA-within, H_3,24_=86.0, p<0.0001; *post hoc*, p<0.0001]. The NKS-3 condition shows the highest within cortex connectivity [VEH *vs*. NKS-3-within, p<0.0001], while the NKS-5 condition shows a similar-to-VEH within cortex connectivity [VEH *vs*. NKS-5-within, p=0.061]. However, both conditions show a significantly reduced inter-community score [VEH *vs*. NKS-3-inter, p<0.0001; VEH *vs*. NKS-5-inter, p=0.0006] (see **Figure 3L,P**). Basal ganglia show a moderate within- and inter-community connectivity in the control condition. The LA, NKS-3 and NKS-5 treatments significantly decreased the connectivity score within the basal ganglia community and towards the rest of the network [VEH *vs*. LA-inter, F_3,1184_=93.7, p<0.0001; *post hoc*, p<0.0001; VEH *vs*. LA-within, H_3,24_=106.3, p<0.0001; *post hoc*, p<0.0001; VEH *vs*. NKS-3-inter, p<0.0001; VEH *vs*. NKS-3-within, p=0.0029; VEH *vs*. NKS-5-inter, p<0.0001; VEH *vs*. NKS-5-within, p<0.0001] (**Figure 3M,Q**). The thalamic structures here studied are poorly connected within the community and with the rest of the network in the control condition and only a significant effect of LA and NKS-3 treatments was found, with both significantly reducing their inter-community connectivity [VEH *vs*. LA-inter, F_3,416_=16.4, p<0.0001; *post hoc*, p<0.0001; VEH *vs*. NKS-3-inter, p=0.0014] (**Figure 3N,R**). Finally, hindbrain’s structures in the control condition are moderately connected to the network, but highly connected to themselves. The three treatments showed a significant effect by reducing both inter- and within-hindbrain community connectivity [VEH *vs*. LA-inter, F_3,656_=54.1, p<0.0001; *post hoc*, p<0.0001; VEH *vs*. LA-within, H_2,24_=35.0, p<0.0001; *post hoc*, p<0.0001; VEH *vs*. NKS-3-inter, p<0.0001; VEH *vs*. NKS-3-within, p<0.0001; VEH *vs*. NKS-5-inter, p=0.0002; VEH *vs*. NKS-5-within, p=0.058] (**Figure 3O,S**).

## Discussion

Fat-taste enhancers, i.e. NKS-3 and NKS-5, targeting CD36 and GPR120 receptors have been developed aiming at *in vivo* modulating human innate preference for dietary fat. They have been, therefore, conceived as a pharmacological tool able to influence oro-sensory detection of dietary lipids in a broader way of lipid sensing, and to alter fat-eating behavior in the mid-long run [14]. The findings of this preclinical study show that prolonged exposure to NKS-3 and NKS-5 induces a functional reorganization of the brain reward network without altering emotional behavior in physiological conditions. In accord with our previous results reporting the absence of aversive responses in a conditioned taste aversion task [14], the fat-taste enhancers did not change how mice behaved in a battery of tests sensitive to the emergence of anxious, depressive, amotivational and compulsive behaviors. This is a key finding considering that NKS-3 and NKS-5, when acutely administrated, induce respectively a 142 and 95 more potent attractive response than natural long-chain fatty acids [14], and no effect of their sustained action neither on their consummatory nor in their emotional behavior has been observed.

As a drinking supplement, a prolonged exposure to the fat-taste enhancers, which derived from the molecular structure of LA, induced a functional reorganization of the brain reward network. This reorganization mostly consisted in a rearrangement of the coordination centers, with for example, fewer participation of the hindbrain’s structures. Most importantly, NKS-3 and NKS-5 affected the organization of the reward network without compromising its stability and the theoretical efficiency of its connections. This finding is particularly obvious while comparing to the effect of LA supplementation. We previously showed that acute lingual application of LA activates brain regions within the reward system [7], and more recently, that long-term LA supplementation does not alter emotional behavior in adult male mice [16]. The unstructured and fragmented reward network obtained in the present study might reflect the loss of the gustatory properties of LA as it was presented at a high concentration, and fatty acids are known to see their hedonic effects reduced or inversed from palatable to unpleasant at high concentrations [26]. Functional network analyses have been proven useful to reveal altered patterns of brain connectivity across different pathological populations, including from neurological and psychiatric contexts (see [27] for review). In our analysis, we exposed the topology of strong and significant connections within the reward network in each experimental condition by filtering weak links representative of spurious connections. That allowed us to evaluate the treatments’ effects on network functional segregation and integration through the identification of nodes and links, i.e. the effects on the ability to process restricted gustatory information within a dense and interconnected brain system. The network reorganization consequence of NKS-3 and NKS-5 exposure reflects the adaptation of the reward system to the gustatory flow information originated in CD36 and GPR120 activation. The analyses revealed that high inter- and within-community scores shared in cortical and hindbrain structures in the control network where affected by the treatments, suggesting that these hubs may play a key role in mediating the treatments’ effects through the modulation of neuronal activity on disparate parts of the reward system. The analyses support that this reorganization does not compromise the global efficiency of the system. However, complementary studies are required in order to confirm these effects over extended and continual exposure, as well as in female individuals. Notwithstanding, we speculate that beneficial effects of fat-taste enhancers on weight control are mediated through the modulation of the reward system activity, against fat preference, via the creation of an artificial satiety state specific for the taste modality. Particularly interesting are the results showing an absence of behavioral perturbations and toxic effects [14] of the treatments, together with an unaltered activity of the basal ganglia within the reward system. Activation of the basal ganglia circuits, which are key players in the transition towards consummatory addictive behavior, influences reward-driven and motivated behavior, as well as movement and motor patterns [28]. Since emotionality and consummatory behavior towards food and beverage were unaffected, the sum of results supports our hypothesis of a slight risk of the agents in terms of addictive potential. Non-negligible is, however, the significantly decreased neural activity observed in the LC of NKS-5-treated mice. Even though the role of the LC in homeostatic feeding is not yet clear, preclinical data have reported the suppression of noradrenergic LC activity during feeding in a satiety state dependent manner [29]. These data are therefore consistent with our hypothesis of fat-taste enhancers as pharmacological tools to induce specific fat satiety. Notwithstanding, we choose to stay cautious in our interpretation of the pharmacological effects on functional connectivity, especially since brain networks remain mathematical representations of a complex real biological system.

It is of common consensus that feeding regulation relies on two parallel systems, the homeostatic and the brain reward systems, which interact to influence food intake [30]. While the former hormonally regulates hunger, satiety and adiposity levels in order to control the energy balance (see [31] for review), the reward system modulates the motivational effects mediated by palatable food, sometimes overriding homeostatic signaling. How the reward system influences food intake is not yet fully understood, but it is essential to take its role into consideration in order to develop new pharmacological tools for treating pathologies as obesity. Future preclinical research including a female population will certainly shed light into the involvement of the reward system in the behavioral and feeding response to long-term exposure to fat-taste enhancers NKS-3 and NKS-5 in the context of obesity. These investigations are to be key in the evaluation of the suitability of these new pharmacological agents as an alternative therapeutic strategy of weight control.

## Acknowledgements

The authors thank the Animal Facilities of Besançon for the technical support and PhD Pierre-Yves Risold (LINC) for his helpful insight.

## Authors contribution

Conceptualization of the experimental design: LC, NK, VVW. Methodology, investigation and data acquisition: LC, LJ, CH, EH, AH. Data analysis and interpretation: LC and VVW. Drafting of the original manuscript: LC and VVW. Supervision: VVW. All authors critically revised the work and approved the version to be published.

## Data availability statement

The data presented in this study are available on request from the corresponding author.

